# Re-emergence of BTV serotype 4 in North Macedonia, July 2020

**DOI:** 10.1101/2020.08.20.259457

**Authors:** John Flannery, Simon King, Paulina Rajko-Nenow, Zagorka Popova, Kiril Krstevski, Igor Djadjovski, Carrie Batten

## Abstract

Bluetongue virus serotype 4 (BTV-4) was confirmed in sheep in North Macedonia in July 2020. The full genome of this BTV-4 strain (MKD2020/06) was shown to be most closely related (99.74% nt identity) to the Greek GRE2014/08 and the Hungarian HUN1014 strains, indicating the re-emergence of this BTV serotype in the Balkan region since it was last reported in 2017.

---

Bluetongue (BT) is a haemorrhagic disease of ruminants which is caused by the BT virus (BTV), the type species of the genus *Orbivirus* within the *Reoviridae (Maclachlan, 2009)*. BTV is typically transmitted by heamatophagous *Culicoides* species which can cause the disease to spread vast distances (Carpenter et al., 2009). There are 24 classical serotypes of BTV (BTV-1 to BTV-24) which are notifiable to the World Organization for Animal Health (OIE) (Kundlacz et al., 2019). In June 2014, a new BTV-4 strain was detected in southern Greece and BT spread throughout the Balkan Peninsula in what was then a BTV-4 naïve ruminant population. The highest morbidity (8.5%) and case fatality rates (37%) were observed in sheep, followed by goats and cattle (Kyriakis et al., 2015) although the severity of illness was not consistent throughout the Balkan countries (Katsoulos et al., 2016). The BTV-4 outbreak extended beyond the Balkan countries to Austria in 2016, and France in 2017 (OIE). The last recorded outbreak of BTV-4 from the Balkan countries was reported to the OIE in 2017 with an outbreak occurring in the Zlatborski region of Serbia.

In July 2020, Macedonian veterinarian authorities notified the OIE of BT in sheep in the Makedonski Brod and Kicevo municipalities. Farmers notified veterinary authorities of suspected BT in sheep on their small holdings, which were located in the same region as the initial report of BT in 2014. On the 8^th^ July, an official veterinarian visited 3 holdings and found that the sheep were exhibiting classical signs of BT such as facial oedema, fever, lameness and recumbency. Twelve EDTA blood samples were taken and were sent to the Faculty of Veterinary Medicine in Skopje who detected BTV-4 using RT-qPCR and partial sequence analysis of BTV segment 2. EDTA blood samples (n=6) from the three holdings were submitted to the OIE reference laboratory for BT at The Pirbright Institute, UK, for confirmatory diagnosis. Nucleic acid was extracted from EDTA blood samples using the LSI MagVet Universal isolation kit (ThermoFisher Scientific) on the KingFisher Flex (ThermoFisher), and BTV was detected using an RT-qPCR assay targeting BTV segment 10 (Hofmann et al., 2008). BTV serotype identification was performed using an assay targeting BTV segment 2 (Maan et al., 2016) for serotype BTV-1, BTV-4, BTV-8 and BTV-9. BTV was isolated on *Culicoides sonorensis* KC cells, following which full genome sequencing was performed using the Nextera XT DNA Library Prep kit (Illumina) on the Illumina MiSeq platform. Sequences were assembled using the BWA-MEM tool (Li and Durbin, 2010) as previously described (Rajko-Nenow et al., 2020), and a neighbor joining tree of BTV segment 2 was built using the default settings in MEGA7.0 software (Kumar et al., 2016). The full genome sequence of isolate MKD2020/06 has been deposited in GenBank under accession numbers MT879201 to MT879210.

All samples (n = 6) were positive using the segment-10 RT-qPCR assay (C_T_ range 18.75–23.10) and using the BTV-4 serotype-specific RT-qPCR assay confirming the clinical diagnosis and the re-emergence of this serotype in the Balkan region since it was last reported in 2018. No amplification was observed for the segment-2 targeted RT-qPCR assays for BTV-1, BTV-8 or BTV-9. A phylogenetic analysis of segment 2 showed the Macedonian 2020 isolate (MKD2020/06) was most closely related to BTV-4 field strains isolated during the previous Balkan BTV-4 epizootic (Figure1). Pairwise alignments of full genomes were performed on EMBOSS Needle (https://www.ebi.ac.uk/Tools/psa/emboss_needle/) using default settings. Across all 10-segments, MKD2020/06 shared 99.74% nucleotide (nt) identity with BTV-4 field strains from Hungary (HUN2014) and Greece (GRE2014/08) in 2014. A BLAST analysis of BTV segment 2, the determinant of BTV serotype, against the available BTV-4 segment-2 sequences revealed that MKD2020/06 was most closely related to: BTV-4/17-15 (99.76%), HUN2014 (99.66%), GRE2014/08 (99.73%) and BTV-4/16-03 (99.56%). The sequences of the remaining segments were also subjected to a BLAST analysis to find the closest relative(s) (Table 1). Based on the %nt identity and the identification of the closest relative in individual segments, MKD2020/06 is a re-emergent BTV-4 strain rather than a novel reassortant virus.

**Figure 1.**
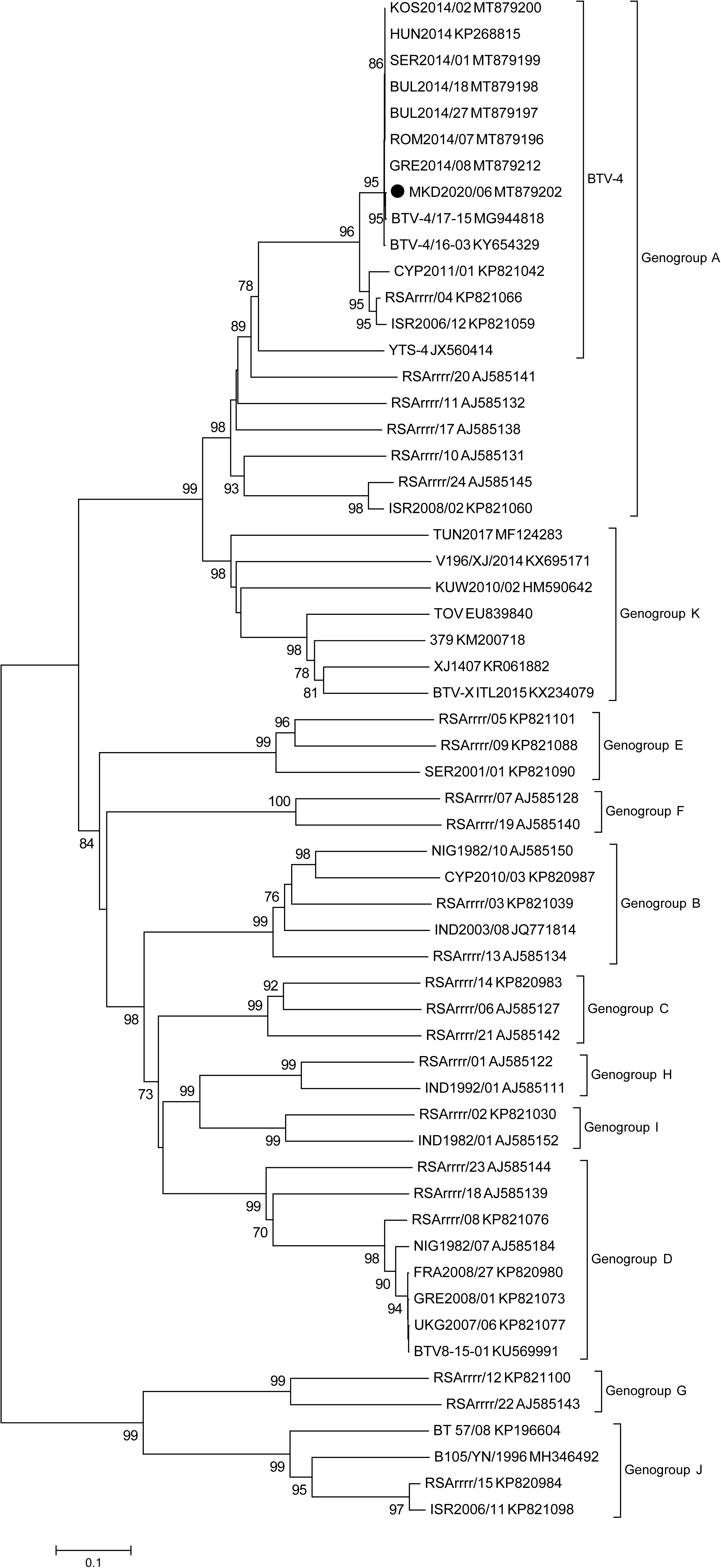
Phylogenetic relationship of BTV-4 (MKD2020/06) from an outbreak in Northern Macedonia in 2020 (identified by black circle) to publicly-available BTV isolates based on segment 2 of the genome. A neighbour-joining tree (1,000 bootstrap replicates) was constructed using the MEGA7.0 software.

**Table 1.** The percent of nucleotide sequence (%nt) identity for individual segments of BTV-MKD2020/06 with the closest matching BTV sequences in GenBank.

| Segment | Length (bp) | GenBank accession no. | Strain ID | %nt identity |
| --- | --- | --- | --- | --- |
| 1 | 3944 | MG944817 | BTV-4/17-15 | 99.82 |
|  |  | KP268814 | HUN2014 | 99.72 |
|  |  | KY654328 | BTV-4/16-03 | 99.70 |
|  |  | MT879211 | GRE2014/08 | 99.77 |
| 2 | 2926 | MG944818 | BTV-4/17-15 | 99.76* |
|  |  | KP268815 | HUN2014 | 99.66 |
|  |  | KY654329 | BTV-4/16-03 | 99.56 |
|  |  | MT879212 | GRE2014/08 | 99.73 |
| 3 | 2772 | MG944819 | BTV-4/17-15 | 99.89* |
|  |  | KP268816 | HUN2014 | 99.82 |
|  |  | KY654330 | BTV-4/16-03 | 99.78 |
|  |  | MT879213 | GRE2014/08 | 99.86 |
| 4 | 1981 | MG944820 | BTV-4/17-15 | 99.85 |
|  |  | KP268817 | HUN2014 | 99.90 |
|  |  | KY654331 | BTV-4/16-03 | 99.70 |
|  |  | MT879214 | GRE2014/08 | 99.90 |
| 5 | 1774 | MG944821 | BTV-4/17-15 | 99.83 |
|  |  | KP268818 | HUN2014 | 99.89 |
|  |  | KY654332 | BTV-4/16-03 | 99.89 |
|  |  | MT879215 | GRE2014/08 | 99.89* |
| 6 | 1637 | MG944822 | BTV-4/17-15 | 99.57* |
|  |  | KP268819 | HUN2014 | 99.39 |
|  |  | KY654333 | BTV-4/16-03 | 99.45 |
|  |  | MT879216 | GRE2014/08 | 99.39 |
| 7 | 1156 | MG944823 | BTV-4/17-15 | 99.83 |
|  |  | KP268820 | HUN2014 | 100 |
|  |  | KY654334 | BTV-4/16-03 | 99.91 |
|  |  | MT879217 | GRE2014/08 | 99.65* |
| 8 | 1125 | MG944824 | BTV-4/17-15 | 99.73* |
|  |  | KP268821 | HUN2014 | 99.73 |
|  |  | KY654335 | BTV-4/16-03 | 99.64 |
|  |  | MT879218 | GRE2014/08 | 99.73 |
| 9 | 1049 | MG944825 | BTV-4/17-15 | 99.70* |
|  |  | KP268822 | HUN2014 | 99.62 |
|  |  | KY654336 | BTV-4/16-03 | 99.52* |
|  |  | MT879219 | GRE2014/08 | 99.62 |
| 10 | 822 | MG944826 | BTV-4/17-15 | 99.88 |
|  |  | KP268823 | HUN2014 | 99.64 |
|  |  | KY654337 | BTV-4/16-03 | 99.64 |
|  |  | MT879220 | GRE2014/08 | 99.51 |

\*Query cover <100%

Small ruminant production has historically been an important part of the Macedonian economy. Approximately 44% of the population live in rural regions with farming (typically in smallholdings) providing a dominant part of the agriculture sector (Food and Agriculture Organization of the United Nations, 2020). Indeed, in 2019 there were approximately 685,000 sheep and 87,500 goats (Republic of North Macedonia State Statistical Office, 2020). Transboundary diseases such as BT present a significant concern as they endanger the sustainability of farming and ultimately food security. While the Macedonian BTV-4 strain has not yet been associated with mortalities in sheep, concomitant infections can exacerbate BT in the field (Kyriakis et al., 2015) thus this BTV-4 strain may yet have an impact on the health status of sheep in affected flocks. The reemergence of BTV-4 should be of concern to the Balkan and neighbouring countries as the possibility exists that it may take the course of the previous epizootic in 2014 whereby 6 countries were affected within 4 months of the outbreak with significant animal health impacts and trade restrictions.

## Acknowledgments

This research was funded by the Department for Environment, Food and Rural Affairs, grant number SE26081 and the Biotechnology and Biological Sciences Research Council, through projects BBS/E/I/00007036 and BBS/E/I/00007037. The authors would like to thank Dr Marc Guimera for supplying cells used for virus isolation.

## Ethical approval

This investigation was performed at the Pirbright Institute in their capacity as an OIE-Reference laboratory for BT and as such, no ethical approval was required for the work carried out.

## Data availability statement

The data that support the findings of this study have been deposited in GenBank under accession numbers MT879201 - MT879210 (MKD2020/06 segment 1 to 10), MT879211 - MT879220 (GRE2014/08 segment 1 to 10), MT879196 (ROM2014/07 Segment 2), MT879197 (BUL2014/27 Segment 2), MT879198 (BUL2014/18 Segment 2), MT879199 (SER2014/01 Segment 2), MT879200 (KOS2014/02 Segment 2).

